# Multiparametric Extracellular Vesicle Profiling Identifies a Distinct Circulating Signature of Bicuspid Aortic Valve Disease

**DOI:** 10.64898/2026.11.03.763956

**Authors:** Luisa Weiss, Amarri Harrison, Sarvesh Chelvanambi, Taku Kasai, Mandy E. Turner, Mark C. Blaser, Marie Billaud, Masanori Aikawa, Sasha A. Singh, Elena Aikawa

**Author notes:** Correspondence: **Luisa Weiss**, PhD Center for Interdisciplinary Cardiovascular Sciences, Brigham and Women’s Hospital, Harvard Medical School, 3 Blackfan Street, Boston, MA 02115, USA School of Biomolecular and Biomedical Sciences, University College Dublin, Dublin, Ireland, **Elena Aikawa**, MD, PhD Center for Interdisciplinary Cardiovascular Sciences, Brigham and Women’s Hospital, Harvard Medical School, 3 Blackfan Street, Boston, MA 02115, USA.

## Abstract

**Background:** Calcific aortic valve disease (CAVD) resulting in life-threatening aortic stenosis, heart failure and death, develops earlier and progresses more rapidly in patients with bicuspid aortic valves (BAV) than tricuspid aortic valves (TAV), yet the circulating molecular drivers remain poorly understood. Extracellular vesicles (EVs) capture disease-relevant cellular activation and offer a powerful window into a BAV-specific pathophysiology.

**Methods:** Circulating large extracellular vesicle (LEV) abundance and morphology were characterized by complementary single-vesicle nano-flow cytometry and super-resolution microscopy. Proteomic profiling further delineated LEV molecular cargoes and associated biological pathways between patients with BAV and TAV.

**Results:** We demonstrate that patients with BAVs exhibited fundamentally altered circulating LEV profiles compared to TAVs. Super-resolution imaging revealed BAV-specific organization of platelet- and leukocyte-associated adhesion molecules into discrete EV membrane microdomains. Proteomic profiling identified distinct LEV cargoes between BAV and TAV, including a BAV-specific, previously unrecognized neutrophil-associated signature enriched for proteins involved in neutrophil degranulation and innate immunity. Consistent with this phenotype, BAV displayed alterations in platelet- and neutrophil-associated LEV populations, suggesting enhanced (thrombo-)inflammatory activity. Lastly, distinct associations between LEV populations and clinical parameters in BAV and TAV suggest that LEV profiles reflect morphology-specific aspects of disease severity.

**Conclusions:** Comprehensive single-vesicle analysis and bulk proteomic profiling identified a distinct circulating LEV signature in BAV-associated CAVD, characterized by endothelial activation and reciprocal platelet-neutrophil interactions. These findings reveal previously unrecognized differences in circulating LEVs between BAV and TAV and highlight their potential as biomarkers of valve morphology-specific pathophysiology.

## 1. Introduction

Aortic valve stenosis (AS) due to calcific aortic valve disease (CAVD) is the most common valvular heart disease in high-income countries and represents a major cause of cardiovascular morbidity and mortality. Bicuspid aortic valves (BAV), the most prevalent congenital cardiac malformation with a prevalence of 1% in the general population, is the strongest non-modifiable risk factor for accelerated AS, with affected individuals typically requiring intervention one to two decades earlier than patients with physiological tricuspid aortic valves (TAV).^1^ Despite increasing evidence that BAVs and TAVs represent pathobiologically distinct forms of AS, comparative mechanistic studies have largely been confined to valve tissue and resident cells. While differential extracellular matrix architecture^2^ and calcification patterns^3^, and distinguish BAVs and TAVs at the tissue level, advances in multi-omics and spatial profiling have only characterized valve cell phenotypes^4^ and valve morphology-and disease stage-specific molecular programs in TAV.^5,6^ In either instance, whether these pathobiological differences are also reflected systemically remains largely unknown.

Congenital valve morphology alone cannot explain the increased susceptibility of BAV to calcification. The fusion of two cusps fundamentally alters valve biomechanics, generating asymmetric blood flow patterns, elevated wall shear stress at the ascending aorta and low magnitude oscillatory shear at the valve cusps.^7^ These hemodynamic disturbances activate mechanosensitive cells, such as valve endothelial cells (VECs) and circulating platelets. Situated at the blood–valve interface, VECs directly sense and respond to the mechanical forces generated by local blood flow.^8^ Endothelial dysfunction may be particularly relevant in BAVs, which exhibit impaired endothelial function, altered nitric oxide signaling and increased oxidative stress in aortic aneurysmal tissue^9–11^, potentially reflecting the combined influence of congenital and hemodynamic abnormalities.

Accelerated and turbulent flow across stenotic valves combined with endothelial disruption exposing thrombogenic surfaces further promote platelet adhesion and activation.^12^ Indeed, activated platelets adhere to endothelial-denuded regions of calcified human aortic valve tissue, and *in vitro*, promote VIC osteogenic transition and mineralization^13^, while platelet-released proteins are associated with accelerated AS progression.^14,15^ Endothelial injury and platelet activation may therefore represent interconnected consequences of pathological valve hemodynamics while simultaneously contributing to continued disease progression.

One mechanism through which endothelial cells, platelets and other activated cells may communicate these pathological signals is the release of extracellular vesicles (EVs). EVs are membrane-enclosed particles that transport proteins, lipids and nucleic acids between cells and can therefore both reflect the phenotype of their cell of origin and modify recipient-cell function.^16^ EVs have emerged as important mediators of cardiovascular calcification^17–20^ and are increasingly implicated in AS, with both valve-derived and circulating EVs carrying inflammatory and osteogenic signals capable of modulating valve mineralization.^21,22^ By providing an interface between the diseased valve tissue and circulation, EVs may reflect cellular activation while simultaneously facilitating intercellular communication. Although EVs from stenotic valve tissue^20,23,24^ and circulating EV-associated miRNAs have been investigated^25,26^, and platelet-and endothelial-derived EV populations increase in patients with aortic stenosis^27^, comprehensive comparisons of circulating EV populations between diseased BAV and TAV are lacking. Thus, it remains unclear whether BAV and TAV generate distinct circulating EV profiles, which cellular populations drive these differences, and how morphological and molecular changes in EVs relate to disease pathophysiology.

Given the distinct hemodynamic environment in BAVs, with associated endothelial dysfunction, platelet activation, and inflammatory signaling, we hypothesized that BAV exhibit a unique circulating EV signature that reflects distinctive molecular processes underlying BAV-associated. Defining these differences may provide key insights into BAV-specific disease mechanisms while identifying potential biomarkers of disease phenotype and progression. The present study reports the first comparative multiparameter characterization of circulating EVs in patients with BAV and TAV. Combining single-vesicle nano-flow cytometry with super resolution microscopy and bulk EV proteomic profiling, we identify marked differences in circulating large EV subpopulations and their molecular cargoes, revealing a previously unreported EV signature associated with neutrophil activation in BAV.

## 2. Methods

### 2.1 Patient Recruitment and Blood Sampling

Patients undergoing surgical valve replacement at Brigham and Women’s Hospital were prospectively recruited. Ethical approval was granted by the Institutional Review Board (2021P001077, 2024P003587, and 2025P001420). All participants provided informed written consent according to the Declaration of Helsinki. Demographic and clinical data were collected including patient age, sex, body mass index, smoking history, valve parameters, medical comorbidities, white cell and platelet count, estimated glomerular filtration rate (eGFR), and concomitant medication (**Supplementary Table 1**). Exclusion criteria included active malignancy, individuals aged <18 years and active or a history of endocarditis. Fasting blood samples were collected from the arterial line into 3x10 mL acid citrate dextrose A vacutainers prior to administration of any sedatives on the morning of valve replacement surgery. Whole blood was sequentially centrifuged as before to generate platelet poor plasma (PPP).^28^ PPP was aliquoted to prevent freeze-thaw cycles and stored at -80°C until further use.

### 2.2 Nanoparticle Tracking Analysis

NTA was performed using ZetaView. Plasma samples were diluted 1:100,000 in 0.1 μm filtered water to a final volume of 1 mL, resulting in an acquisition concentration of 140-200 particles/frame. The manufacturer’s default software settings for EVs were selected. For each measurement, 11 cell positions and 60 frames per position were captured. After capture, the videos were analyzed by the in-build ZetaView Software 8.02 and particle concentration, size and zeta potential were calculated.

### 2.3 EV enrichment

Large EVs were enriched by benchtop centrifugation as before.^29,30^ In brief, 100 μL plasma was centrifuged for 10 min at 3,000 xg and 4°C to pellet cell debris. LEVs were pelleted for 20 min at 20,000 xg and 4°C, washed with 500 μL PBS and pelleted again. The resulting pellet was used immediately for flow cytometry experiments or stored at -20°C for proteomics and immunoblotting.

### 2.4 Large EV Flow Cytometry

Phenotyping of circulating large plasma EVs was performed as described before^29^ according to the MIFlowCytEV framework (**Supplementary Table 2**). The cellular origin was determined by nano-flow cytometry (Cytoflex S, Beckman Coulter) using cell type-specific surface antigens for platelet CD41-PE (303706, BioLegend), CD42b-APC (303912, BioLegend), and CD62P-AF488 (304916, BioLegend), endothelial CD62E-FITC (303114, BioLegend) and CD31-AF594 (303126, BioLegend), as well as neutrophil CD11b-APC (379906, BioLegend) and CD66b (305104, BioLegend) activation. Large EVs were identified based on violet side scatter and Exobrite^TM^ 400 True EV Membrane (30136-T, Biotium) labeling. Representative dot plots detailing the gating strategy are shown in **Figure S1**.

### 2.5 Super Resolution Microscopy

Samples were prepared using the EV Profiler 2 kit (900-00201, Oxford Nanoimaging (ONI)) with phosphatidyl serine (PS) capture. In brief, EVs from three biological donors per cohort were pooled and concentrated for 30 min at 14,000 xg using Amicon® Ultra-0.5 Centrifugal Filter Devices with a 3 kDa cutoff (UFC500396, Millipore), achieving an approximate concentration factor of 10. The EV chip was prepared according to the manufacturer’s instructions. EVs were supplemented with the PS capture supplement, added to the chip and incubated for 2 hours at room temperature. Custom antibodies for detection were CD41-Alexa Fluor 647 (303726), CD62P-Alexa Fluor 488 (304916), CD31-Alexa Fluor 594 (303126), CD62E-Alexa Fluor 488 (BBA16G, R&D Systems), CD11b-Alexa Fluor 488 (393107), CD66b-Alexa Fluor 647 (392911). All antibodies were purchased from BioLegend unless otherwise indicated. All antibodies were used at a concentration of 10 μg/mL. Cell Mask Orange (C10045, ThermoFisher Scientific) was used at a concentration of 50 ng/mL. 12 random positions per lane were acquired. Super-resolution images of EVs were acquired using dSTORM equipped with a 100x, 1.4NA oil immersion objective. The temperature was set at 30°C and the illumination angle for TIRF imaging was 53°. Before capturing images, fluorescent channels were calibrated for colocalization using the bead slide provided by the manufacturer. Colocalization analysis on single EVs was performed with the CODI online analysis platform (https://alto.codi.bio/) using the AutoEV application and clustering was performed using the AI Essentials EV Profiling Model 3.

To assess whether the relationship between EV cluster area and the number of localizations differed between valve morphologies, linear mixed-effects models were fitted separately for each EV population. Number of localizations and cluster area were log10-transformed, and models included cluster area, valve morphology (BAV versus TAV), and their interaction as fixed effects, with field of view (FOV) included as a random intercept to account for clustering of observations within individual FOVs. The area-valve morphology interaction was used to determine whether the scaling relationship between cluster area and the number of localizations differed between BAV and TAV. Spearman correlation coefficients were calculated separately by valve morphology for visualization of within-group associations.

### 2.6 ELISA

ELISA kits to quantify plasma levels of MPO (DMYE00B, Quantikine) and neutrophil elastase (DY9167-05, Duoset) were purchased from R&D Systems. The MPO ELISA was performed according to the manufacturer’s instructions. As the neutrophil elastase ELISA is a commercially available development kit, titration of the detection antibody and the Streptavidin-HRP reagent was necessary. The optimized assay utilized the detection antibody at 62.5 ng/mL and a 13.3-fold dilution of the Streptavidin-HRP reagent.

### 2.7 Mass spectrometry sample preparation

5 ug protein, determined by BCA, were used as input. Mass spectrometry sample preparation was performed using the PreOmics iST-BCT kit (P.O.00099) according to the manufacturer’s instructions. In brief, samples were lysed, reduced, and alkylated for 10 min at 95 °C and 1000 rpm, transferred to a cartridge and subsequently double-digested with LysC and trypsin at 37 °C and 500 rpm for 1 hour. Samples were washed twice with provided wash buffers and eluted using the provided elution buffer. Samples were evaporated at 45 °C and peptides resuspended in LC-load buffer to a final peptide concentration of 100 ng/µL.

Tryptic peptides were analyzed using a quadrupole Orbitrap Exploris 480 coupled to a Vanquish Neo UHPLC system (Thermo Fisher Scientific) as before. 400 ng peptides were fractionated using a dual column setup: a PepMap Neo C18 Trap Cartridge, 5 μm, 300 μm × 5 mm (Thermo Fisher Scientific); and an Easy-Spray PepMap Neo C18 Column, 2 μm, 75 μm × 150 mm (Thermo Fisher Scientific). The column was heated at a constant temperature of 45 °C. The gradient flow rate was 300 nL/min from 5 to 21% solvent B (95% acetonitrile/0.1% formic acid in mass spectrometry-grade water) for 20 min, 21–30% solvent B for 10 min (total acquisition, 30 min), and another 15 min of a 95–5% solvent B sawtooth wash. Solvent A was 0.1% formic acid in water. The mass spectrometry analysis was performed by using data-independent acquisition (DIA). MS1 scans were acquired in profile mode, and MS2 scans were acquired in centroid mode. Spray voltage was set to 2000 V, funnel radio frequency (RF) level at 50, and heated capillary temperature at 275 °C. MS1 was with 120 K resolution, and mass scan range was set to m/z 400–900. The AGC target was set to standard with a maximum injection time of 25 ms. MS2 spectra were acquired with a precursor isolation range of m/z 400–900 divided into 12 Th windows with an overlap of 1 Da. MS2 resolution was set to 15 K and the AGC target value for fragment spectra was set at 1000% with a maximum injection time of auto. Higher-energy collisional dissociation (HCD) collision energy was set to 26%.

### 2.8 Mass spectral annotation and data availability

The spectra were queried against the UniProt human database (downloaded May 2025; 42,421 entries). Trypsin/P was set as the digestion enzyme, allowing up to 2 missed cleavages and a minimum peptide length of 7–30 amino acids (1–6 peptide charge). Oxidation (+15.995 Da) of methionine was set as a variable modification, and carbamidomethylation (+57.021 Da) of cysteine was set as a static modification. The Quantification Type was set to the default setting MS2 Apex (Quan in all files). All files were exported in one combined consensus proteome. Proteins quantified in at least 3 donors in at least one cohort and with at least 1 unique peptide (1% FDR) were included in the statistical analysis.

The mass spectrometry proteomics data have been deposited to the ProteomeXchange Consortium via the PRIDE partner repository with the dataset identifier PXD084233.

### 2.9 Immunoblotting

For orthogonal validation of proteins differentially expressed by mass spectrometry, 10 μg protein per donor was resuspended 6x reducing Laemmli Buffer (BP-111R, Boston BioProducts) and boiled at 95°C for 10 min. For assessment of neutrophil granule proteins, LEVs were further concentrated as described above before sample preparation for gel electrophoresis. Proteins were separated on a 4-20% Mini-PROTEAN TGX Precast Gel (456-1096, BioRad) prior to protein transfer onto a polyvinylidene fluoride membrane. Membranes were blocked in 5% skim milk for 1 hour at room temperature before incubation with primary antibodies at a dilution of 1:500 overnight at 4 °C. Monoclonal antibodies used include CD62P (84298T, Cell Signaling Technologies), EMILIN1 (NBP-84127-25, Novus Biologicals), Osteopontin (ab8448, Abcam), Thrombospondin 4 (MAB2390, R&D Systems), Myeloperoxidase (ab25989, Abcam), Neutrophil Elastase (ab131260, Abcam), Ficolin-1 (ab325599, Abcam), CD58 (330902, Biolegend, 1:250), and Cathepsin D (ab75852, Abcam). Membranes were then incubated with goat anti-rabbit HRP (7074S, Cell Signaling Technologies) or goat anti-mouse HRP (31430, Thermo Fisher Scientific) secondary antibodies for 2 hours at room temperature and imaged using enhanced chemiluminescence substrate in an Amersham ImageQuant 800. Signal intensities were quantified using ImageJ and normalized to the total protein amount (Albumin).

### 2.10 Statistical Analysis

All statistical analyses and data visualization were performed in RStudio (version 2024.9.0.375).

Differences in EV abundance between BAV and TAV derived from nano-flow cytometry and NTA, and protein abundance determined by ELISA and densitometry were assessed using two-sided Wilcoxon rank-sum tests. *p*<0.05 was considered statistically significant. Cluster-level metrics derived from super-resolution microscopy were compared between BAV and TAV using two-sided Wilcoxon rank-sum tests, performed separately for each EV population and cluster metric. The relationship between cluster area and the number of molecular localizations was assessed using linear mixed-effects models. Both cluster area and number of localizations were log10-transformed prior to analysis. Models included area, valve morphology (BAV or TAV), and their interaction as fixed effects, with field of view (FOV) included as a random intercept to account for clustering of observations within each FOV. The interaction term was used to determine whether the relationship between cluster area and number of localizations differed between BAV and TAV. Analyses were performed separately for each marker-defined EV population, and interaction *p*-values were adjusted for multiple comparisons using the Benjamini–Hochberg method. Data were visualized using linear regression lines with 95% confidence intervals.

For proteomics analysis, differential protein abundances between BAV and TAV were assessed using linear modeling with empirical Bayes moderation of protein-wise variances implemented in the limma package. Statistical significance was determined from the moderated *t*-statistics, with multiple-testing correction applied using the Benjamini–Hochberg false discovery rate (FDR) at 0.05.

Functional enrichment analysis was performed using Enrichr-KG.^32^ Gene Ontology (GO) and Reactome pathway databases were queried to identify significantly over-represented pathways. Enrichment significance was assessed using Enrichr-derived *p*-values, with multiple-testing correction.

## 3. Results

### Distinct large EV populations are elevated in BAV

While previous studies have reported increased levels of in circulating large EVs (LEVs) in AS compared to healthy controls^27^ or changes following transcatheter aortic valve replacement^32–34^, comparative characterization of EV subpopulations and their proteomic cargoes between BAV and TAV patients has never been performed. We hypothesized that the congenital hemodynamic differences in BAV are reflected by increased activation of mechanosensitive cells, such as platelets and endothelial cells. To identify shared and differing molecular signatures of circulating EVs, we combined single-vesicle analysis by nano-flow cytometry and super resolution microscopy with label-free quantification proteomic characterization of circulating plasma EVs. 24 patients undergoing valve replacement surgery for severe aortic stenosis (n=10 TAV, n=14 BAV) were prospectively recruited at Brigham and Women’s Hospital. Patient cohorts were well matched for age, sex, comorbidities and hemodynamic indices (**Supplementary Table 1**).

Fasting blood samples were collected on the morning of surgery for plasma EV analysis. Characterization of small plasma EVs (sEVs) <200 nm by nanoparticle tracking analysis revealed no difference in sEV concentration, size and zeta potential between the groups (**Figure S2**). These results may not be surprising, given that shear stress, particularly oscillatory shear or disturbed flow, cause rapid plasma membrane remodeling, primarily inducing LEV release through membrane blebbing.^35,36^ We then employed our previously published method to enrich plasma for LEVs by benchtop centrifugation (**Figure 1A**)^28^. Nano-flow cytometry indicated a nearly two-fold increase in the total concentration of LEVs in BAV PPP (*p*=0.00085, **Figure 1B**). We further investigated whether *in vivo* activation resulted in a shift in specific LEV subpopulations using a panel of well-defined CD markers (**Figure 1C**). We observed an increase in circulating CD42b-positive (*p*=0.032) and CD62P-positive (*p*=0.0081) EVs in BAV while platelet-derived CD41-positive EVs remained unaltered. Moreover, CD31-positive (*p*=0.00084) and CD62E-positive (*p*=0.0022) LEVs were significantly increased in BAV (**Figure 1D**). Since expression of E-selectin (CD62E) is a selective marker of inflammatory and cytokine-activated endothelial cells, these results may indicate aberrant endothelial activation in BAV. As CD62E can mediate neutrophil recruitment to vascular walls through CD11b^37^, we also measured neutrophil-derived EVs. C66b-positive EVs were significantly higher (*p*=0.00051), while the concentration of CD11b-positive EVs only indicated a trend towards an increase in BAV patients (*p*=0.07). Since CD11b is constitutively expressed on the neutrophil and macrophage membrane, while CD66b externalizes from internal granules during neutrophil activation and degranulation, these findings may indicate heightened neutrophil activation in BAV.

**Figure 1:**
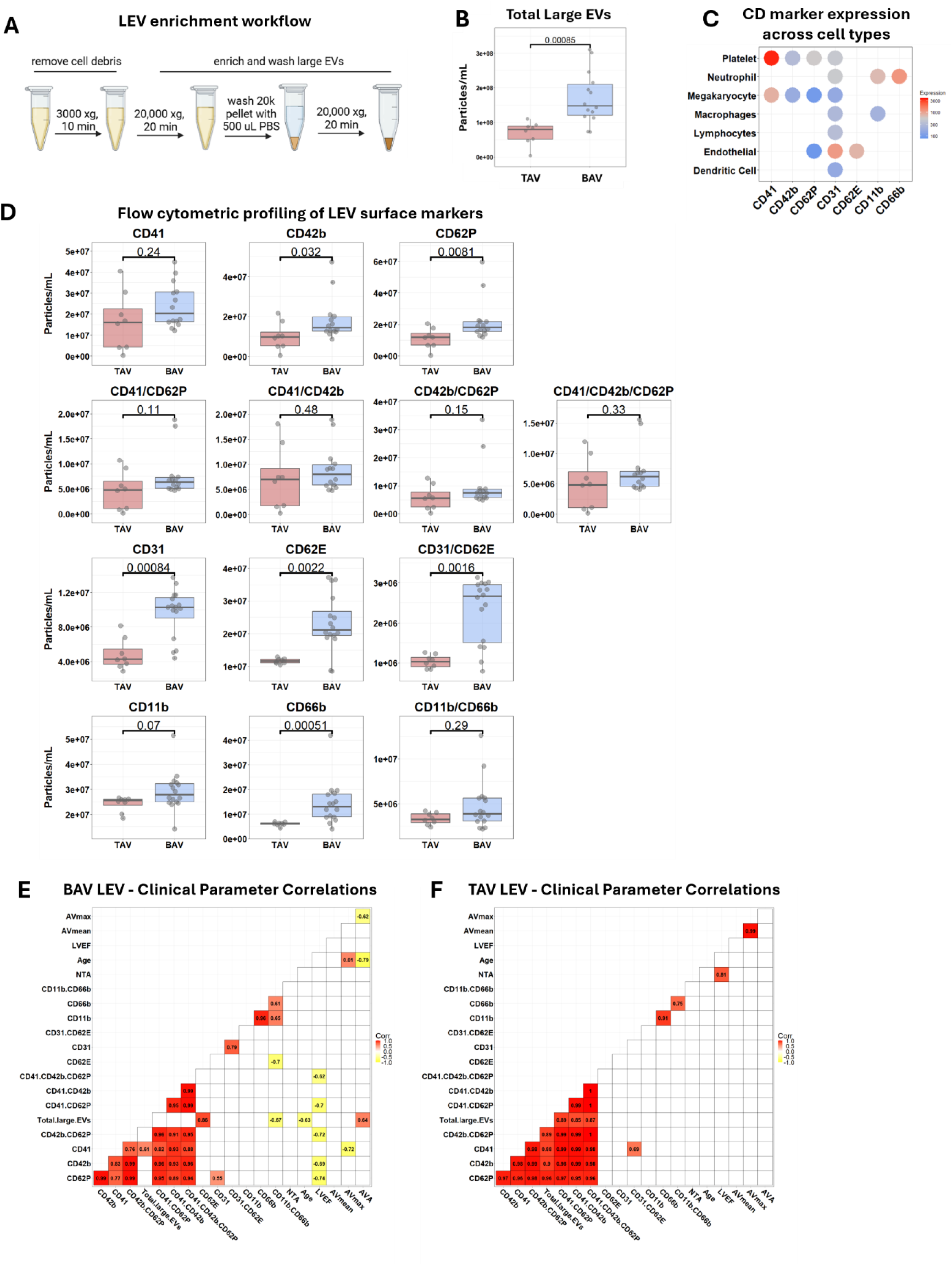
Distinct large EV populations are elevated in BAV. (A) Schematic overview of LEV enrichment by benchtop centrifugation. (B) Total LEV levels quantified by nano-flow cytometry. Statistical significance was determined using a two-sided Wilcoxon ranksum test. Data are presented as box-and-whisker plots, with the box indicating the interquartile range (IQR), the center line representing the median, and whiskers extending to 1.5× IQR. Individual points represent individual donors. (C) CD marker expression (normalized counts per million, nCPM) across cell types was derived from single-cell data from the Human Proteome Atlas. (D) Nano-flow cytometry profiling of the cellular origin of LEVs. Platelet-derived EVs (CD41, CD42b, CD62P, CD41/CD62P, CD41/CD42b, CD42b/CD62P, and CD41/CD42b/CD62P), endothelial EVs (CD31, CD62E, and CD31/CD62E) and leukocyte/neutrophil EVs (CD11b, CD66b, and CD11b/CD66b) were quantified. BAV (n=14), TAV (n=10). Statistical significance was determined using a two-sided Wilcoxon rank sum test. Data are presented as box-and-whisker plots, with the box indicating the interquartile range (IQR), the center line representing the median, and whiskers extending to 1.5× IQR. Individual points represent individual donors. (E) Spearman correlation analysis of clinical parameters with LEV counts in BAV and (E) TAV. Significant (p<0.05) correlations are shaded, and correlation coefficients are denoted.

To further assess the clinical relevance of variations in circulating LEV populations, we examined associations between LEV concentrations and clinical parameters. Stratification by valve morphology revealed distinct relationships between LEV concentration and clinical parameters in BAV (**Figure 1E**) and TAV (**Figure 1F**). Notably, several (primarily platelet-derived) LEV subpopulations inversely correlated with left ventricular ejection fraction (LVEF) exclusively in BAV (**Figure 1E**), indicating that higher circulating platelet-derived LEV levels associate with impaired left ventricular systolic function. Collectively, these findings demonstrate distinct circulating LEV profiles in BAV and TAV, with prominent endothelial and neutrophil activation in BAV AS patients.

### Platelet and neutrophil EVs exhibit distinct membrane microdomain organization in BAV

To gain deeper insight into the morphology and molecular characteristics of the LEVs, we performed super resolution microscopy. LEVs bleb off the plasma membrane in response to cellular activation, a process associated with the externalization of phosphatidylserine (PS), a procoagulant phospholipid highly implicated in cardiovascular disease. Therefore, PS-based capture was employed to assess PS-positive LEVs independently of their tetraspanin surface composition. For detection of all captured LEVs, irrespective of their membrane marker expression, CellMask Orange (CMO) was used as a general LEV stain. In line with our nano-flow cytometry results, the total number of EVs, quantified as detected clusters per field of view (FOV, **Figure 2A**), was increased in BAV (*p*=0.0042), which also translated into an increased number of total EV clusters detected (**Figure S3A**). Moreover, LEVs in BAV display a larger mean size (*p*=0.026, **Figure 2B, D**; **Supplementary Table 3**) with a concomitant decrease in circularity (*p*<0.0001, **Figure 2C, D**; **Supplementary Table 3**), potentially signifying a more complex EV composition.^38^ We further investigated these effects on cell-specific LEV populations. Corroborating our nano-flow cytometry results, the number of CD62P-, CD31-, CD31/CD62E-, and CD66b-positive LEVs were increased in BAV (**Figure S3B**). Conversely, CD42b and CD62E-positive LEV levels did not change, and we observed an increase in CD11b- and CD11b/CD66b-positive LEVs, previously undetected by flow cytometry (**Figure S3B**). Since LEVs for super resolution microscopy were captured based on their PS exposure, these EVs represent a subpopulation of the overall EVs quantified by flow cytometry. Together, these findings suggest that BAV is associated with a distinct enrichment of neutrophil-derived, PS-exposing LEVs, revealing phenotypic differences not captured by conventional nano-flow cytometry.

**Figure 2:**
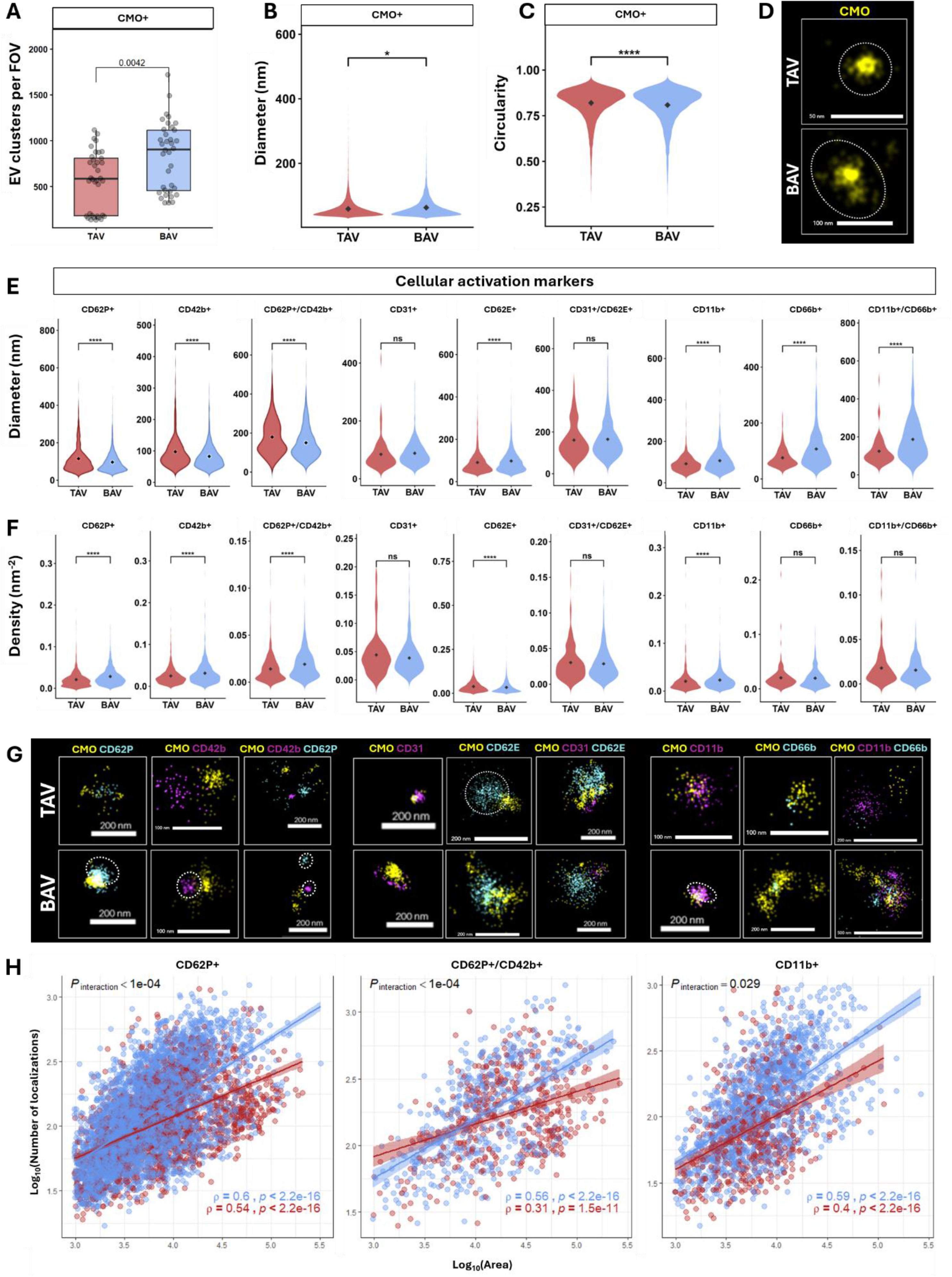
Super Resolution Microscopy reveals distinct membrane microdomain organization in BAV LEVs. (A) Average number of quantified EV clusters of CellMask Orange (CMO)-positive EVs per field of view (FOV, n=36 per group). Data are presented as box-and-whisker plots, with the box indicating the interquartile range (IQR), the center line representing the median, and whiskers extending to 1.5×IQR. Individual points represent individual FOVs. (B) Quantification of the mean diameter (nm) of CMO-positive EV clusters. Statistical significance was determined using a two-sided Wilcoxon rank sum test followed by Benjamini Hochberg correction for multiple comparisons. Data are presented as violin plots, with the black diamond indicating the mean. (C) Quantification of the mean circularity of CMO-positive EV clusters. Statistical significance was determined using a two-sided Wilcoxon rank sum test followed by Benjamini Hochberg correction for multiple comparisons. Data are presented as violin plots, with the black diamond indicating the mean. Full statistical analysis of all measured parameters including n-numbers (Supplementary Table 3). (D) Representative images of CMO-positive EVs in TAV (top) and BAV (bottom) sample. Scale bars provided for each panel. Dashed area’s approximate size and shape of the respective EV cluster. (E) Quantification of the mean diameter (nm) and (F) mean density (/nm2) of platelet (CD42b, CD62P, and CD42b/CD62P), endothelial (CD31, CD62E, and CD31/CD62E), and neutrophil (CD11b, CD66b, and CD11b/CD66b) EV clusters. Statistical significance was determined using a two-sided Wilcoxon rank sum test followed by Benjamini Hochberg correction for multiple comparisons. Data are presented as violin plots, with the black diamond indicating the mean. Full statistical analysis of all measured parameters including n-numbers (Supplementary Table 3). (G) Representative images of CMO-positive EVs in TAV (top) and BAV (bottom) sample. Scale bars provided for each panel. Circled areas indicate dense localizations. (H) Scatter plots show log_10_-transformed cluster area and number of localizations for marker-defined EV populations in BAV and TAV. Individual points represent individual clusters. Lines represent linear regression with 95% confidence intervals. Differences in the relationship between cluster area and number of localizations by valve morphology were assessed using linear mixed-effects models incorporating an area×valve interaction and field of view as a random intercept. Displayed *P*_interaction_ values are BH-adjusted.

Further harnessing the power of super resolution microscopy, we quantified the morphological characteristics of our LEV populations at single-vesicle level. Individual EVs from spatially grouped single-molecule fluorescence localizations are reconstructed by the EV profiling model, with each resulting localization cluster representing a detected EV. Single EV cluster-level metrics, including the number of localizations, area, diameter, circularity, and localization density (**Supplementary Table 3**), are subsequently derived from the spatial distribution and geometry of the localizations within each cluster, enabling quantitative assessment of EV size, morphology, and marker organization. While CD42b- and CD62P-positive EVs exhibited a smaller mean diameter in BAV, CD11b-, CD66b-, and CD62E-positive EVs were significantly larger compared with TAV (**Figure 2E**; **Supplementary Table 3**). Despite their smaller size and similar number of localizations (**Supplementary Table 3**), BAV LEVs displayed tight and spatially distinct clustering of CD62P and CD42b on the EV surface (**Figure 2F and G**). Larger clusters furthermore contained proportionally more molecular localizations in BAV than in TAV, particularly within CD62P-positive, CD62P/CD42b-double positive, and CD11b-positive clusters (**Figure 2H**), suggesting altered organization of marker-positive molecular domains in BAV EVs. Consistent with the increased density of localizations (**Figure 2F and G**), these results indicate differential spatial organization of marker-positive domains between valve morphologies.

### BAV LEVs exhibit a distinct neutrophil-associated proteomic signature

Given the markedly altered LEV profiles identified by nano-flow cytometry and super resolution microscopy, we next sought to establish whether these differences were also reflected in the proteomic composition of enriched LEVs. To directly link differences in proteomic cargo to the LEV subsets characterized above, we applied the same LEV enrichment strategy (**Figure 1A**). Mass spectrometry-based proteomic profiling quantified 915 LEV proteins, with inter-donor correlations of *r*>0.89 and an average between-group correlation of *r*=0.953 (**Figure S4**). Comparison of the identified proteins to previously published data sets of LEVs revealed that 494 of the quantified proteins have previously been associated with LEV from platelets^39,40^, endothelial cells^41^, neutrophils^42^, erythrocytes^43^ and/or plasma lipoproteins^44^ (**Figure 3A**; **Supplementary Table 4**). In contrast to small EVs, for which several canonical marker proteins have been proposed, no consensus constitutive markers are currently established for LEVs, hence Gene Ontology (GO) enrichment analysis was performed as a complementary strategy to provide functional annotation of the complete proteomic dataset and evaluate its association with EV-related cellular compartments. Proteins in our plasma LEV proteome were significantly enriched within the GO cellular component terms “Extracellular Vesicles”, “Extracellular Exosome” and several other vesicle-associated terms (**Figure 3B**; **Supplementary Table 5**), providing further evidence of successful EV enrichment. To more deeply characterize the 421 proteins not previously associated with LEVs, we conducted Reactome pathway enrichment analysis, revealing predominant enrichment in pathways related to the term “Neutrophil Degranulation” and “Innate Immune System” (**Figure S5; Supplementary Table 6**), suggesting that these previously uncharacterized LEV-associated proteins reflect innate immune and neutrophil-related processes. Moreover, these proteins were significantly annotated to “Heart Disease”, “Cardiovascular Disease”, and “Chronic Kidney Disease” in the Jensen Diseases database (**Figure S5**; **Supplementary Table 6**), potentially implicating a clinical relevance of circulating EVs as surrogate disease signatures.

**Figure 3:**
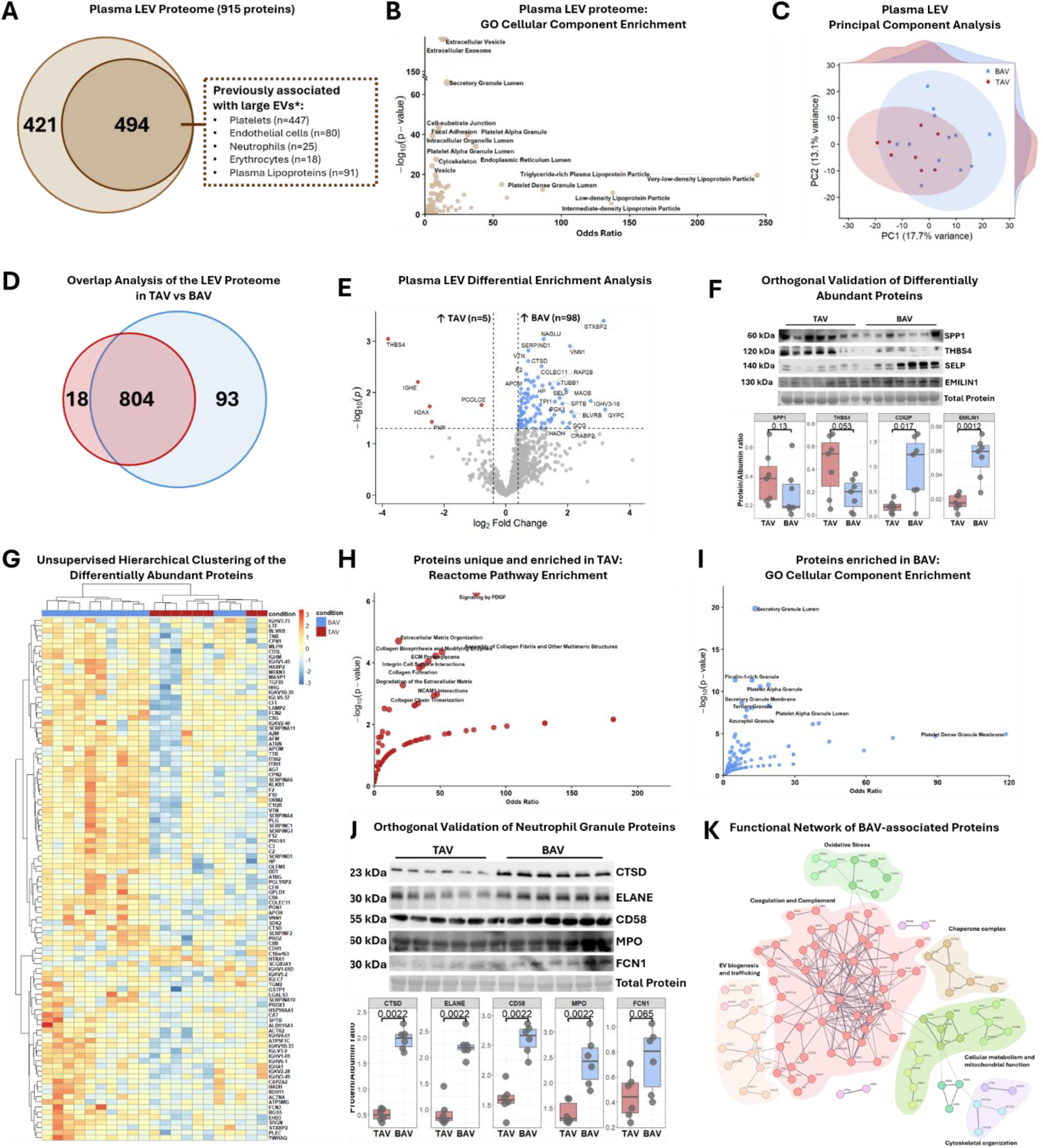
BAV LEVs exhibit a distinct neutrophil-associated proteomic signature. (A) Mass spectrometry-based proteomics of LEVs from BAV (n=12) and TAV (n=8). (A) Venn diagram of quantified proteins and the proportion of proteins previously identified in LEVs from platelets, endothelial cells, neutrophils, erythrocytes, and lipoproteins. [*] indicates that some proteins were identified in multiple data sets. Proteome overlap analysis (Supplementary Table 4). (B) Gene Ontology (GO) Cellular Component enrichment analysis of all quantified LEV proteins. All Cellular Component pathways and annotations (Supplementary Table 5). (C) Plasma LEV principal component analysis with density plots. (D) Venn diagram of the proteins unique to TAV (n=18) or BAV (n=93) or shared between the groups (n=804). Full list of unique proteins (Supplementary Table 6). (D) Volcano plot of differentially enrichment analysis between BAV and TAV using a t-test. Proteins highlighted in red are enriched in TAV (n=5), proteins highlighted in blue and enriched in BAV (n=98, p<0.05, Supplementary Table 8). (F) Orthogonal validation of differentially enriched proteins by immunoblotting and densitometric quantification of osteopontin (SPP1), thrombospondin 4 (THBS4), CD62P (SELP) and Emilin 1. Statistical significance of the relative signal intensity normalized to total protein (unmodified membranes, Supplementary Figure 5) was determined using a two-sided Wilcoxon rank sum test. Data are presented as box-and-whisker plots, with the box indicating the interquartile range (IQR), the center line representing the median, and whiskers extending to 1.5×IQR. Individual points represent individual donors (n=7 per group). (G) Unsupervised hierarchical clustering of the differentially enriched proteins (p<0.05). (H) Reactome pathway enrichment analysis of proteins enriched (n=5) or uniquely quantified (n=18) in TAV. All Reactome pathways and annotations (Supplementary Table 9). (I) GO Cellular Component enrichment analysis of proteins enriched (n=98) in BAV. All GO Cellular Component pathways and annotations (Supplementary Table 10). (J) Orthogonal validation of differential abundance of “Ficolin-1 rich Granule” and “Tertiary Granule” associated proteins cathepsin D (CTSD) and CD58 as well as classic neutrophil degranulation markers neutrophil elastase (ELANE), myeloperoxidase (MPO), and Ficolin-1 (FCN1) by immunoblotting and densitometric quantification of relative signal intensities normalized to total protein (unmodified membranes, Supplementary Figure 6) was determined using a two-sided Wilcoxon rank sum test. Data are presented as box-and-whisker plots, with the box indicating IQR, the center line representing the median, and whiskers extending to 1.5×IQR. Individual points represent individual donors (n=6 per group). (K) Functional Network of BAV-associated proteins. Network based on Reactome pathway enrichment among proteins significantly differentially enriched (n=98) or unique (n=93) in BAV. Protein-protein interaction network was produced by mapping the gene list onto STRING (confidence level >0.700) and k-means clustering partitioned the network into distinct functional modules.

Principal component analysis (PCA) revealed similar global proteomic profiles between the TAV and BAV patient cohorts, highlighting their shared disease etiology. However, partial spatial separation in PC1 may suggest morphology-specific molecular features (**Figure 3C**). Indeed, 18 proteins were uniquely found in TAV LEVs (**Figure 3D**; **Supplementary Table 7**), including osteopontin [SSP1], which was previously identified to be increased in calcified human valve tissue from patients with TAV compared to BAV.^45^ This finding emphasizes the potential of LEVs to reflect valve-specific pathological processes and their capability to serve as circulating biomarkers for AS. Overall, differential protein expression analysis revealed relatively modest differences between groups, with 103/915 identified proteins (11%) exhibiting nominally significant differences in abundance (*p*<0.05; **Figure 3E; Supplementary Table 8**). Given the exploratory nature of the analysis, proteins with nominal significance were retained for hypothesis generation and pathway-level interpretation and independently validated by western blotting to confirm disease-specific abundances across the cohorts (**Figure 3F**). Consistent with the overlap observed in the PCA, unsupervised hierarchical clustering did not fully segregate samples by cohort (**Figure 3G**), reflecting the biological heterogeneity within each patient group and the largely shared proteomic landscape, despite the presence of distinct disease-associated molecular features.

To characterize the key differences between BAV and TAV proteomes, we employed GO analysis. Proteins unique to or increased in TAV LEVs were enriched in GO terms related to extracellular matrix organization and degradation (**Figure 3H; Supplementary Table 9**), potentially reflecting systemic biological processes associated with TAV. Proteins increased in BAV, on the other hand, recapitulated our detected neutrophil degranulation signature, specifically secretion from azurophilic, Ficolin-1-rich, and tertiary granules, while also indicating augmented platelet activation (**Figure 3I; Supplementary Table 10**). To further validate this unique neutrophil signature, we assessed the differentially expressed proteins cathepsin D and CD58 by western blotting (**Figure 3J**), alongside the classical neutrophil granule makers myeloperoxidase (MPO), neutrophil elastase, and ficolin 1, which have previously been identified in calcified valve tissue.^46^ Intriguingly, while MPO and neutrophil elastase EV profiles differed in BAV vs TAV (**Figure 3J**), their overall levels in PPP did not change between the groups (**Figure S6**), highlighting this signature is confined to LEVs.

Lastly, STRING protein–protein interaction analysis of increased or uniquely identified proteins in BAV was performed using a high-confidence interaction threshold (confidence score ≥0.700) to infer a functional network of BAV-associated proteins. K-means clustering partitioned the network into distinct functional modules, revealing enrichment of several interconnected biological pathways (**Figure 3K**), spanning coagulation and complement pathways, EV biogenesis, cellular metabolism, and cytoskeletal organization. Importantly, the identification of mitochondrial dysfunction and oxidative stress pathways aligns with the recognized contribution of these processes to BAV pathophysiology.^11^ Collectively, our single vesicle and proteomics data reflect known disease-associated pathways, while uncovering a previously unreported neutrophil activation signature in patients with BAV, suggesting that EVs may act as circulating surrogate biomarkers for the molecular mechanisms underlying the accelerated disease progression in BAV.

## 4. Discussion

Despite distinct pathophysiological features between BAV and TAV, investigations have largely been confined to valvular tissue morphology and cellular alterations, with comparatively little attention given to systemic molecular signatures such as circulating EVs. We conducted the first comprehensive multi-parameter characterization of circulating EVs in human AS, combining single EV analysis by nano-flow cytometry and super resolution microscopy with the first proteomic assessment of LEV cargoes, with the overarching aim of identifying disease-specific signatures in circulation and their potential as minimally invasive biomarkers. Circulating LEVs derived from (activated) platelets, endothelial cells and neutrophils were increased. Beyond surface marker positivity, super resolution microscopy revealed distinct phenotypic alterations in LEVs, pinpointing to the formation of functionally relevant membrane microdomains. Proteomic characterization of LEV cargoes further uncovered a previously unrecognized role of neutrophil degranulation in BAV, potentially reflecting systemic innate immune activation beyond immune cell infiltration into the valve tissue.

EVs are well-established drivers of cardiovascular calcification^19^; however, the contribution of circulating EVs in AS and their differential role in BAV vs TAV remain elusive. Previous investigations have been limited to comparisons between controls and patients diagnosed with aortic stenosis^27^ or pre- and post-transcatheter^32–34,47^ or surgical^48^ valve replacement analyses, persistently excluding or not disclosing the proportion of BAV patients, leaving disease-specific EV signatures between BAV and TAV largely unexplored. Therefore, we sought to characterize the hitherto unknown changes in the molecular profiles of circulating small and large EVs between these cohorts. Small EV profiles did not differ between BAV and TAV, whereas total LEVs, as well as platelet-, activated endothelial cell-, and neutrophil-derived LEVs were increased in BAV. This preferential increase in LEVs may reflect enhanced shear-mediated vesicle release rather than generalized constitutive cellular activation. In line with our findings, increased levels of CD31-positive endothelial-derived LEVs have been reported in BAV compared to TAV, while levels of CD62E- and CD31/CD42b-positive LEVs remained unchanged.^49^ In this study, however, only 15% of the BAV patients had aortic stenosis, whereas all TAV patients were free of overt disease, suggesting that endothelial dysfunction in BAV may in part reflect underlying congenital valve phenotype rather than established AS alone. Upon disease onset, endothelial dysfunction and platelet activation may become amplified in BAV, promoting the release of LEVs from activated platelets and endothelial cells. Moreover, adhesion molecules, including CD62P, CD42b, and CD11b, formed distinct membrane microdomains on BAV-derived LEVs, while these receptors were more diffusely distributed on TAV LEVs. The formation of CD42b membrane microdomains is vital for platelet activation^50^, and reciprocal interactions of CD62P, CD62E, and CD11b mediate bidirectional neutrophil activation, characterized by the complexation of CD11b with CD18 that subsequently mediates platelet adhesion via CD62P and CD42b.^51^ Mimicking biologically relevant microdomains of the parent cell may therefore allow EVs to functionally recapitulate aspects of their parental-cell phenotype and elicit similar downstream responses in recipient cells.

The well-established concept of reciprocal activation of platelets and neutrophils in cardiovascular diseases has recently gained attention as a potential therapeutic target for thrombo-inflammatory diseases.^52^ CD62P-PSGL-1 (P-selectin glycoprotein ligand 1) and CD42b-CD11b/CD18 (Mac-1) interactions mediate platelet-neutrophil adhesion and activation, while EVs may also transfer these adhesion molecules and facilitate bidirectional signaling between both cell types. Crucially, redistribution of CD62P and PSGL-1 into respective membrane microdomains is essential for platelet-neutrophil aggregate formation, neutrophil recruitment^53,54^ and neutrophil-derived oncostatin M mediated CD62P clustering on human endothelial cells.^55^ The organization of these proteins into discrete domains may therefore increase their local avidity and facilitate platelet-neutrophil/neutrophil-endothelial interactions, potentially contributing to inflammatory signaling in BAV. Moreover, binding of PSGL-1 triggered intracellular signaling in neutrophils, activated Mac-1 and enhanced the adhesive activity of P-selectin, thereby strengthening neutrophil adhesion to platelet and endothelial cells^56,57^ and prompted the generation of (CD11b-positive) neutrophil-derived EVs. Internalization of these EVs by platelets was in turn mediated through interaction of neutrophil Mac-1 and platelet CD42b.^57^ Our observed increase in CD62P-positive EVs in BAV may therefore promote neutrophil interactions and subsequent CD11b-positive neutrophil EV release, establishing a feed-forward mechanism that amplifies inflammatory signaling, although functional validation of this hypothesis is warranted. Of note, inflammatory cell infiltration is increased by 122% in BAV compared with TAV^58^ and platelet-leukocyte interactions and thrombo-inflammation have been implicated in AS progression.^14,59,60^ Our findings extend these observations by suggesting that reciprocal platelet-neutrophil, neutrophil-endothelial activation and EV-mediated signaling represent a previously unrecognized component of the heightened inflammatory phenotype in BAV. Intriguingly, CD62P-induced neutrophil activation also promotes the secretion of cathepsin G and neutrophil elastase^61^, indicating a role for CD62P in holistic neutrophil activation. Since neutrophil elastase levels are increased in serum and valve tissue of patients with AS and alvelestat, a selective inhibitor of neutrophil elastase, ameliorated valve thickening in apoE-deficient mice^62^, targeting neutrophil activation in AS may confer therapeutic benefit.

Besides the modulated vesicle signatures, our proteomic analysis independently identified and validated neutrophil degranulation as a previously unreported BAV-specific pathway. The protein cargo of circulating EVs represents a particularly unexplored dimension of AS. Proteomics provides an unbiased approach to simultaneously characterize hundreds of EV-associated proteins, enabling the identification of differences that extend beyond predefined surface markers and revealing coordinated changes in biological pathways and potential mediators of intercellular communication. Proteomic and multi-omics profiling of valve tissue enabled the discovery of disease stage- and spatially restricted molecular networks not resolved by targeted measurements alone.^5,6^ Similarly, proteomic analysis of tissue-entrapped EVs revealed disease-associated EV proteins and identified previously unrecognized procalcific signaling networks.^24^ While proteomic differences between BAV- and TAV-derived VICs have recently been examined^63^, isolated VICs may undergo phenotypic changes during culture and therefore may not fully recapitulate their *in vivo* state. Moreover, this approach interrogates resident valve cells rather than circulating vesicular signals. To our knowledge, we present the first report of the circulating large EV proteome and delineate pathophysiologically relevant differences between patients with calcified BAV and TAV.

Accumulating evidence highlights the dichotomous role of neutrophils in sterile chronic inflammation in cardiovascular disease, spanning from exacerbation of atherosclerosis and plaque destabilization to recruitment of endothelial progenitor cells for reendothelialization of stenotic arteries and tissue repair.^64–66^ Tight orchestration of granule release is pivotal for neutrophils to exert maximal effector functions, ensuring that potent host-defense responses are restricted to the site of infection or injury. Hence, packaging of effector molecules into distinct granules and their spatio-temporal release in response to external cues allows for targeted host defense. The classical hierarchy of neutrophil granule mobilization following infection results in the sequential release of tertiary, secondary and lastly primary (azurophilic) granules.^67^ However, sterile inflammation triggers stimulus-dependent release of neutrophil granule subtypes^68^ that expand neutrophils’ diverse cellular effects in disease microenvironments. Neutrophil degranulation in AS is previously unreported, likely attributable to the rapid activation and short half-life of neutrophils. Evidence for a contribution of neutrophils in AS is confined to the presence of neutrophil extracellular traps (NETs) in valve tissue that correlated with AS severity^46^ and the potential of plasma from AS patients to induce NET formation *ex vivo.*^69^ The authors speculated about a potential role of shear stress and neutrophil-endothelial interactions as mechanisms underlying this activation and, indeed, shear stress was recently shown to directly mediate NETosis *in vitro*^70^, providing a clinically relevant link to altered hemodynamics in BAV. Given the temporal differences in neutrophil degranulation and NET formation, NETs observed in calcified valves may represent the stable end stage of neutrophil activation while our in-depth characterization of circulating LEVs delineates the transient degranulation that cannot be captured on a tissue level. Importantly, platelets actively sense their environment, selectively sequester circulating biomolecules such as nucleic acids, proteins and EVs into their granules^71,72^, and release them upon activation^73^, potentially facilitating the preservation and systemic dissemination of otherwise transient signals.

Our study has several limitations, including a modest sample size. A particular strength of our study, however, is that BAV and TAV patients were well matched, minimizing potential clinical confounding. In addition, the cross-sectional design precludes conclusions regarding whether the identified LEV signatures contribute to or result from BAV-associated pathophysiology. Moreover, the PS-based enrichment strategy for super resolution microscopy captures a specific subpopulation of circulating EVs; therefore, our findings of the membrane microdomains in BAV may not extend to the entire EV population. While all our analyses converge towards a BAV-specific neutrophil-associated LEV signature, if this phenotype is replicated in sEVs and the functional consequences remain to be established. In particular, assessment of circulating platelet–neutrophil aggregates could provide functional validation of these signatures. However, as this signature emerged during downstream analyses, all samples had already been processed to PPP, precluding assessment of intact platelet–neutrophil aggregates in the available cohort. Finally, the exploratory associations between LEV populations and clinical parameters require validation in larger, independent cohorts.

Collectively, the identification of a neutrophil-associated LEV signature may have functional relevance to AS progression, as increasing evidence indicates that neutrophil activation and NET formation drive cardiovascular inflammation and calcification. Notably, a recent study demonstrated that pharmacological inhibition of NET formation attenuates inflammation, osteogenic signaling, and valve calcification in experimental mouse AS^74^, highlighting neutrophil-mediated pathways as potentially modifiable contributors to disease progression. Integrated characterization of circulating EV concentration, cellular origin, morphological properties, clinical associations, and proteomic cargoes could reveal how the distinct hemodynamic and cellular environments of BAV and TAV interconnect systemically and identify EV-associated molecular signatures underlying their divergent pathophysiology. Holistically addressing this gap may provide insight into biological processes that cannot be captured by analysis of the excised valve alone.

## 5. Acknowledgements

Luisa Weiss is supported by the European Union’s Horizon Europe Research and Innovation Program under the Marie Skłodowska-Curie Actions (101203749). Elena Aikawa’s research is supported by the National Institutes of Health R01HL174066, the Leducq Foundation PRIMA network (22RAF02), a research grant from Pfizer (GR 1000131), the American Heart Association Strategically Focused Research Network (26SFRNCVHD1592182), and Kowa Company, Japan.

## Declaration

No generative AI has been used to generate data presented in this manuscript.

## Supplementary Materials

### Supplementary Tables

**Supplementary Table 1:** Patient demographics.

**Supplementary Table 2:** MIFlowCyt-EV framework.

**Supplementary Table 3:** Super resolution microscopy EV cluster summary statistics.

**Supplementary Table 4:** Overlap of quantified proteins with previously published LEV proteomes.

**Supplementary Table 5:** Gene Ontology Cellular Component pathways of all quantified proteins.

**Supplementary Table 6:** Reactome and Jensen Diseases pathways of proteins previously not associated with LEVs.

**Supplementary Table 7:** Unique and shared proteins between BAV and TAV.

**Supplementary Table 8:** Differentially enriched proteins between BAV and TAV.

**Supplementary Table 9:** Reactome pathways of proteins unique to or enriched in TAV LEVs.

**Supplementary Table 10:** Cellular Component pathways of proteins enriched in BAV LEVs.

### Supplementary Figure Legends

**Supplementary Figure 1:**
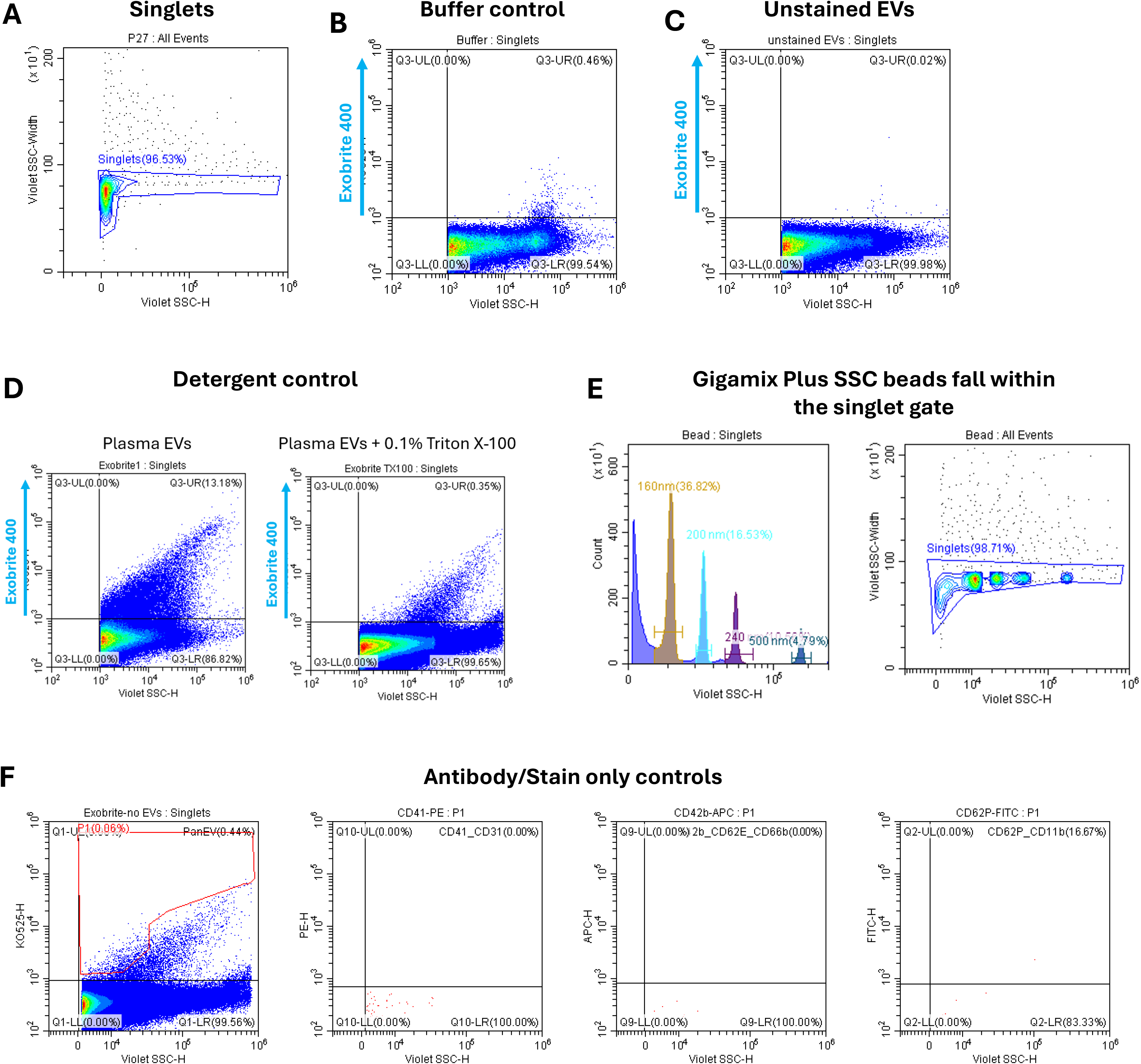
Gating strategy for LEV nano-flow cytometry. (A) Data acquisition was performed on a Beckman Coulter CytoFlex S flow cytometer. (A) Violet Side Scatter (VSSC) Height vs VSSC Width delineates single vesicle gate. (B) A buffer control was analyzed to determine background vesicle counts. (C) Unstained EV were acquired to determine autofluorescence. (D) Exobrite 400 stained EVs were acquired (left) before addition of 0.1% Triton X-100 to the same sample (right). Of the initially 13.18% positive EVs, 0.35% remain after detergent lysis, indicating that acquired particles are indeed EVs. (E) Gigamix Plus SSC beads with defined sizes (160 nm, 200 nm, 240 nm and 500 nm) were used to adjust VSSC intensity (left) and to approximate validate that EVs of these sizes fall within the established single vesicle gate. (F) Antibody/Stain only controls. Exobrite 400 indicates some aggregate formation (left), resulting in generation of gate P1 for vesicle quantification. PE-, APC- and FITC-labelled antibodies do not indicate aggregates when gates for P1.

**Supplementary Figure 2:**
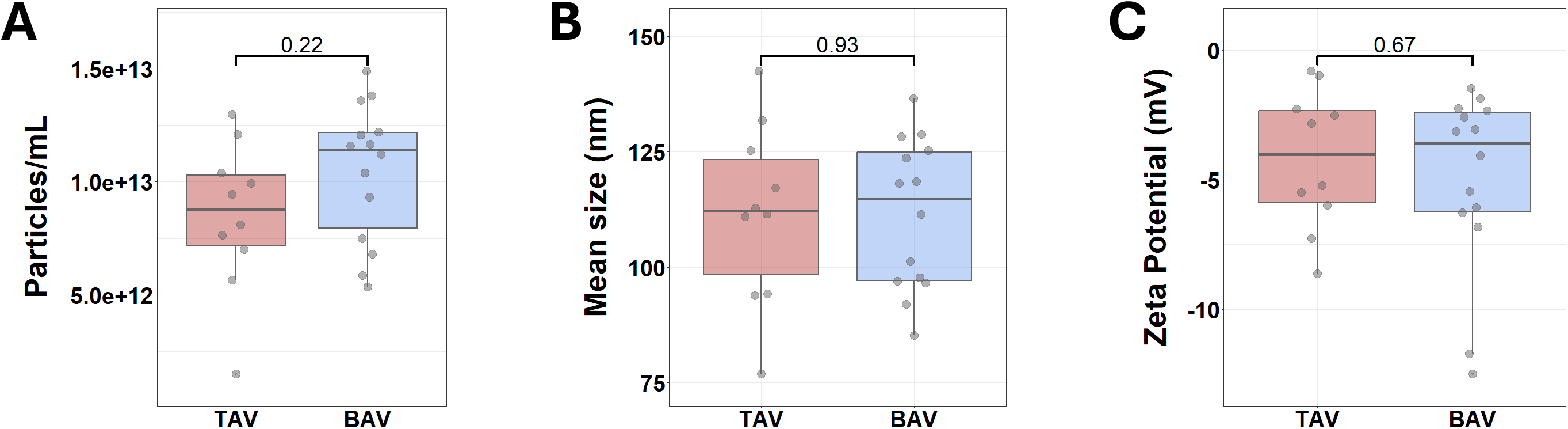
Nanoparticle Tracking Analysis does not indicate differences in small EV levels. (A) Nanoparticle Tracking Analysis was performed using a ZetaView. Plasma samples were diluted 1:100,000 in 0.1 um filtered water, resulting in 140-200 particles/frame for analysis of small EVs (<200nm). (A) Total particles (/mL), (B) mean size (nm) and (C) zeta potential (mV) of small EVs do not differ between the groups. BAV (n=14), TAV (n=10). Statistical significance was determined using a two-sided Wilcoxon rank sum test. Data are presented as box-and-whisker plots, with the box indicating the interquartile range (IQR), the center line representing the median, and whiskers extending to 1.5×IQR. Individual points represent individual donors.

**Supplementary Figure 3:**
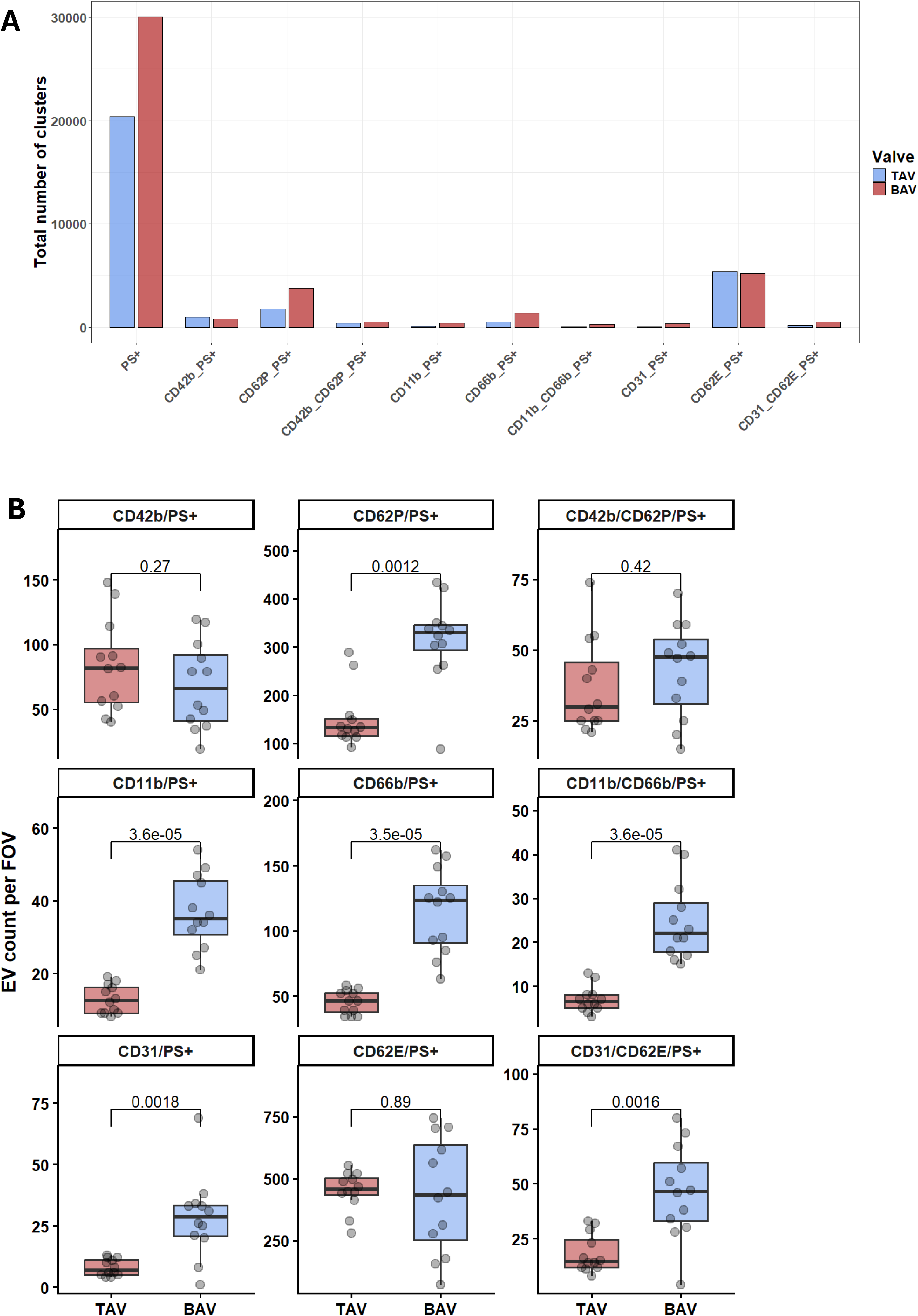
Total number of quantified EV clusters and EV clusters per field of view and per marker. (A) Total number of quantified EV clusters summed up over n=36 FOVs each for CMO-positive EV clusters or n=12 FOVs for all other markers. (B) Average number of quantified platelet (CD42b, CD62P, and CD42b/CD62P), endothelial (CD31, CD62E, and CD31/CD62E), and neutrophil (CD11b, CD66b, and CD11b/CD66b) EV clusters per field of view (FOV, n=12 per group). Statistical significance was determined using a two-sided Wilcoxon rank sum test followed by Benjamini Hochberg correction for multiple comparisons. Data are presented as box-and-whisker plots, with the box indicating the interquartile range (IQR), the center line representing the median, and whiskers extending to 1.5×IQR. Individual points represent individual FOVs.

**Supplementary Figure 4:**
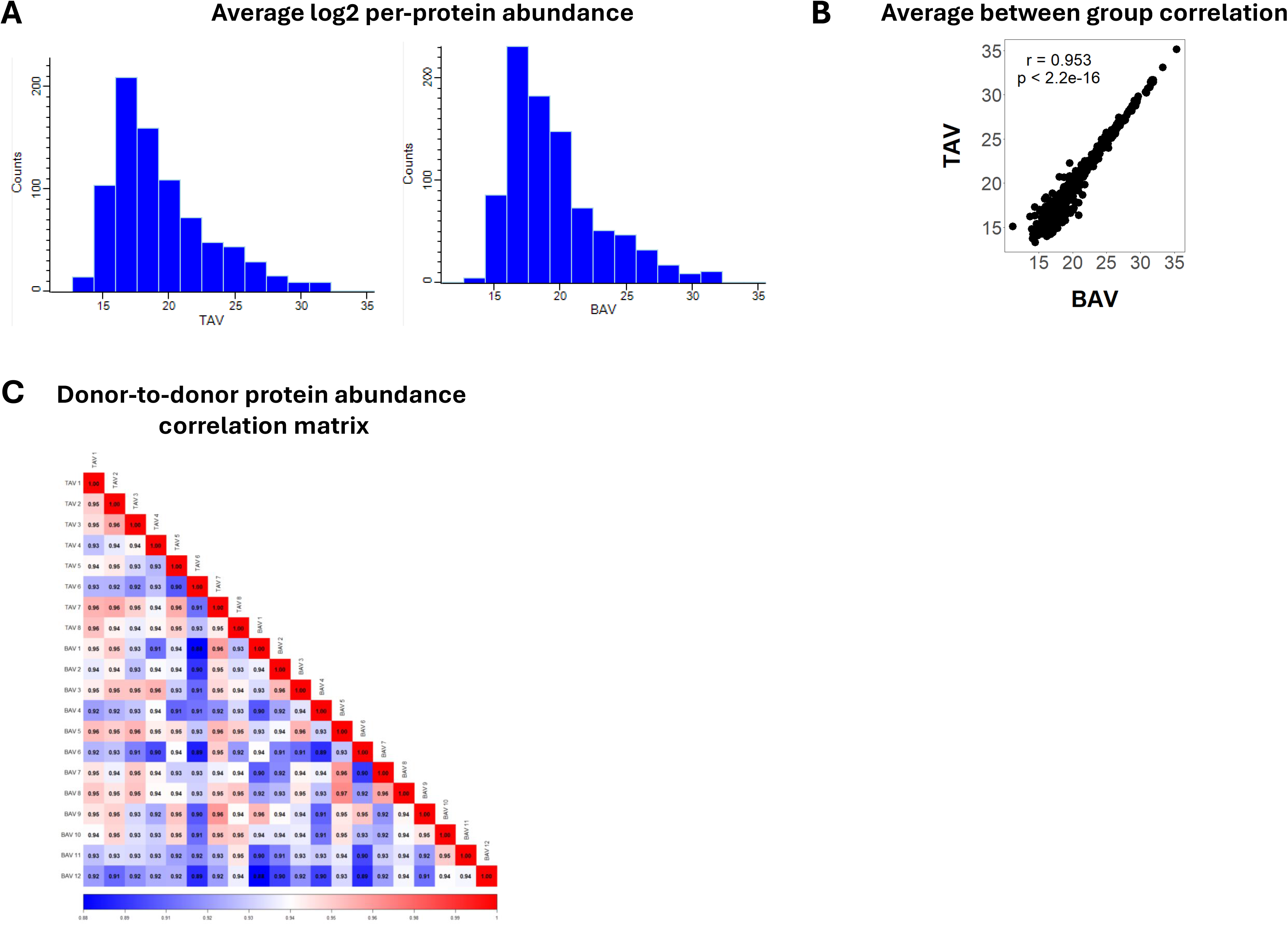
Quality control of proteomics data. (A) Frequency distribution histograms of mean log2 per-protein abundances by sample group (B) Scatter plot of valve-specific mean protein abundances with associated spearman correlation coefficient and *p*-value. (BAV n=12, TAV n=8). (C) Spearman correlation analysis of donor-to-donor protein abundance.

**Supplementary Figure 5:**
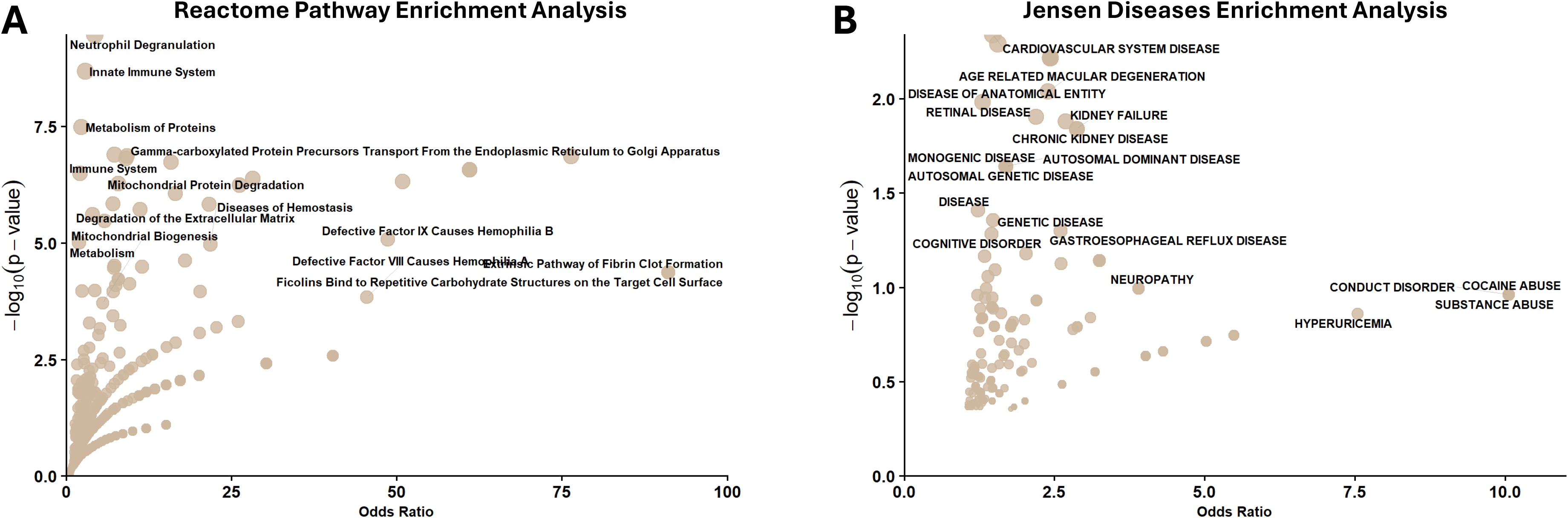
Extended Gene Ontology analysis of proteins previously not associated with LEVs. (A) Reactome pathway enrichment analysis of proteins previously not associated with LEVs (n=421, corresponding to Figure 3A). All Reactome pathways and annotations (Supplementary Table 6). (B) Jensen Diseases enrichment analysis of proteins previously not associated with LEVs (n=421, corresponding to Figure 3A). All Jensen Diseases pathways and annotations (Supplementary Table 6).

**Supplementary Figure 6:**
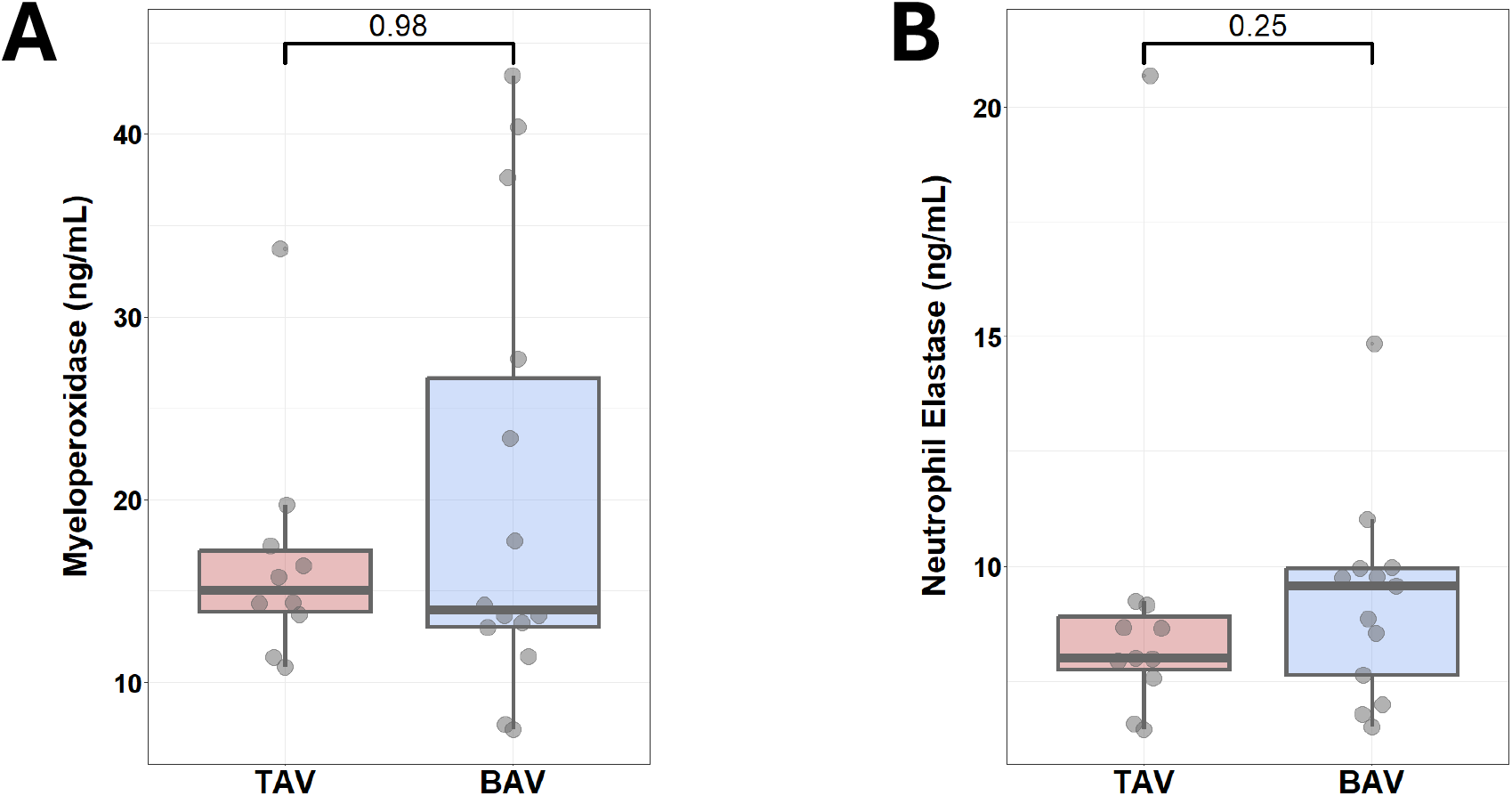
Neutrophil degranulation signature is confined to LEVs. Quantification of (A) MPO and (B) neutrophil elastase plasma levels by ELISA. Statistical significance was determined using a two-sided Wilcoxon rank sum test. Data are presented as box-and-whisker plots, with the box indicating the interquartile range (IQR), the center line representing the median, and whiskers extending to 1.5×IQR. Individual points represent individual donors (TAV n=10, BAV n=14).

**Supplementary Figure 7:**
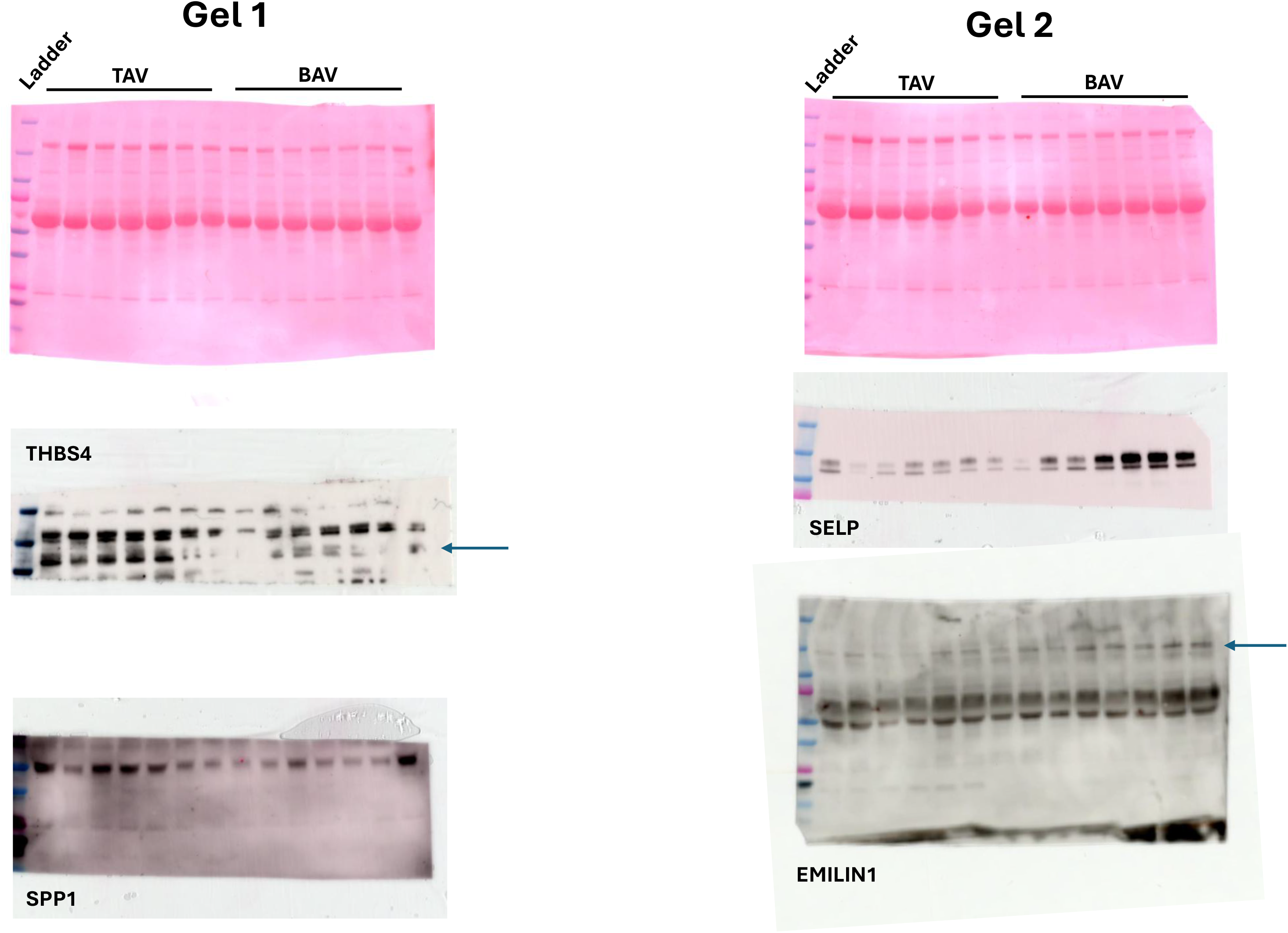
Uncropped immunoblots for Figure 3F.

**Supplementary Figure 8:**
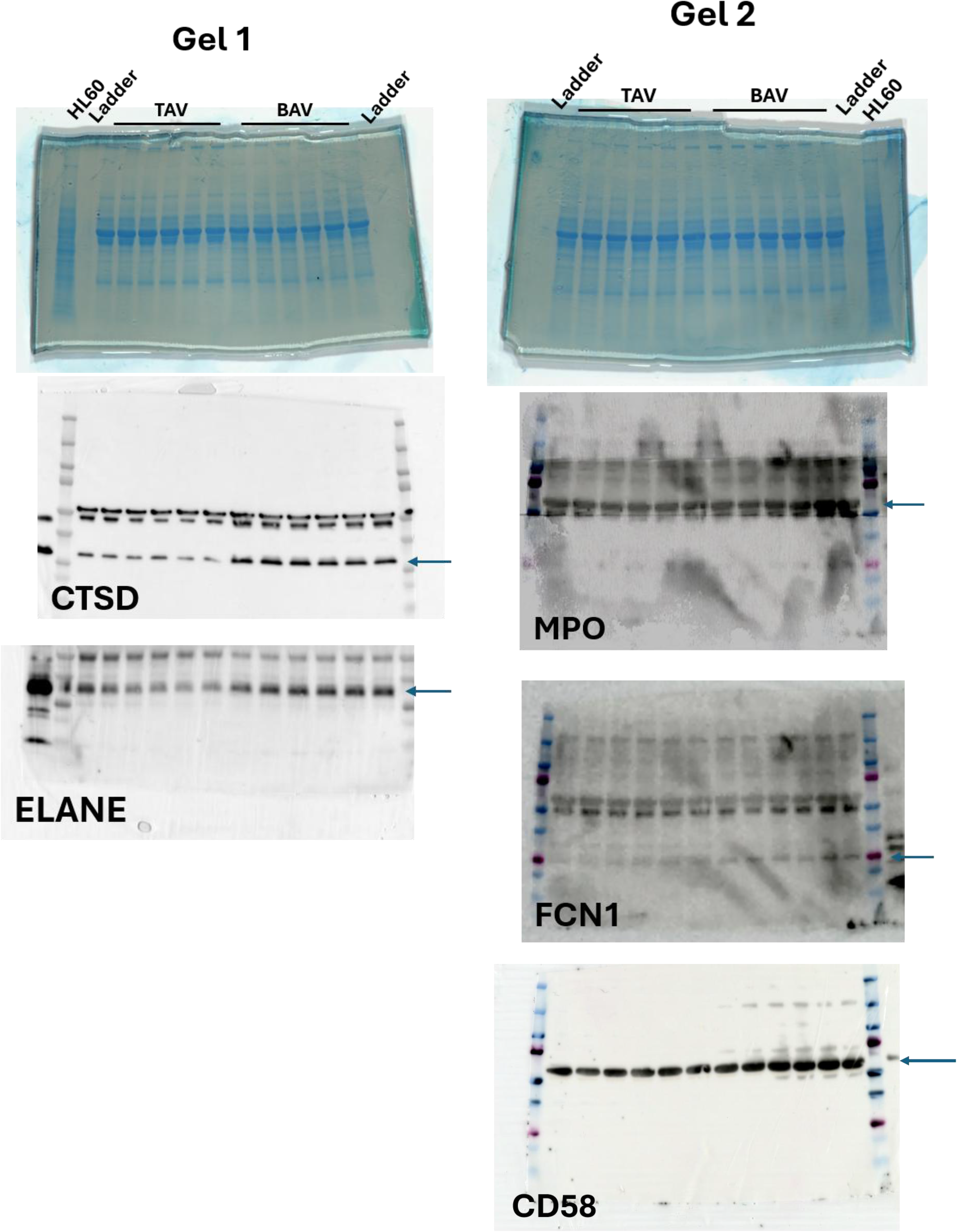
Uncropped immunoblots for Figure 3J.

